## supplementary figure legends and table for "Social buffering switches fear to safety encoding by oxytocin recruitment of central amygdala “buffer neurons”"

**Supplementary material**

**Supplementary Figure 1: Effects of SBF and retention of SBF are immediate, starting from the first representation of the CS.**

Representation of same results as in Fig. 1b, but with the values of freezing responses to the individual 4 consecutive exposures to CS1 that receive SBF on Day 3 and that are separately averaged per presentation. The presence of the companion on Day 2 (SBF) does not gradually decrease freezing to the CS, but leads to an immediate decrease in freezing levels (i.e. with the start of the first CS exposure). On Day 3, the maintenance of this decrease is also immediate (occurring upon exposure to the first CS) and maintained at equal low level throughout subsequent exposures. Mean values ±SEM are shown.

**Supplementary Figure 2: Immediate and long-lasting effects of SBF in animals under different housing conditions or of different sex.**

**(a)** In group-housed animals the presence of the companion acutely and long-lastingly reduces freezing similar to all other previously obtained results. n=8 no companion, n=11 + companion *p<0.05; ***p<0.001, Two way ANOVA, all Bonferroni-corrected (**b**) Animals of different sex exhibit acutely and long-lastingly reduced freezing levels to similar extent in the presence of a companion of the same sex (males: filled bars; females: striped bars). For both (a) and (b), rats were exposed (as described in protocol in Figure 1a) during habituation on Day 1 four times to two different auditory conditioned stimuli (CS1: 5 kHz, red; CS2: 15 kHz, blue) and subsequently fear conditioned by pairing each CS with an electric foot shock of 0.5 mA. On Day 2 memory of fear was assessed by re-exposure to each CS (fear recall). 3 hours later, rats were habituated to a polyester ball (n=6) or a companion rat in the adjacent compartment (n=6) for 10 min and subsequently re-exposed to the CS2 (SBF). On Day 3, both CSs were presented again in the absence of a companion (Retention of SBF). Freezing was diminished only for the CS (CS2) that was paired with social buffering. Conventions as in Fig. 1. Two-way ANOVA with repeated measures (pre-tone, CS1, CS2) and group ("+ No companion, + Companion"), for each session (Habituation, Fear recall, SBF, Retention of SBF). Individual and mean values ± SEM are shown. **(c)** SBF works equally well in the presence of a familiar or unfamiliar conspecific in a contextual fear conditioning paradigm. Two-way ANOVA followed by Tukey correction for multiple comparisons; *, p<0.05; **, p<0.01; blue asterisks, Brother vs Without Companion; green asterisks, Stranger vs Without Companion. n=4 for all groups.

**Supplementary Figure 3 | Analysis of behavioral interactions between fear-conditioned and companion rats during different experimental paradigms.**

(**a**) Bar chart of behavioral activity of demonstrator rat in absence or presence of companion rat before recall of the CS on Day 2 with and without injection of OTA. For these analyses, we assessed the behavior every second (bin-size of ethograms=1 sec) using the following categories: "exploration" ("EX"), "freezing", "grooming" (licking of entire body), "social interaction" (nose to nose or nose to body through the plexiglass wall, "SI"), "social motivation" (close to the plexiglass wall, "SM"). Active behavior was defined as the total time spent in exploration, social interaction and social motivation. Individual and mean values ± SEM are shown. (n=5-7) **(b)** Ethograms of color-coded behaviors before, during and after 4 CS presentations to FC-rats and their companion with each line at similar vertical position in upper and lower blocks representing behavior of resp. an individual FC-rat and its corresponding companion. First series: vehicle injected in CeA of FC-rat (black, n=5), second series: OTA injected in FC-rat (green, n=7), third series: FC-rat exposed to polystyrene ball (grey, n=6). For these analyses, we assessed the behavior every second (bin-size of ethograms=1 sec) **(c)** Bar chart of average time exhibiting social behavior (="social interaction" + "social motivation") by FC-rats injected with vehicle (black) or OTA (green) and their corresponding companions (white). Two-way ANOVA followed by Tukey correction for multiple comparisons; *, $p<0.05$ **(d)** Scatter plot of social behavior levels (as defined in (C) as a function of freezing levels in FC-rats injected with vehicle (black) or OTA (green).

**Supplementary Figure 4: Chemo,- and optogenetic effects are specific to the viral expression.**

**(a)** IP CNO does not disrupt SBF in the absence of DREADD expression. CNO, given IP 30 min before SBF does not affect decreased freezing levels during "SBF" and these decreased freezing levels again appear during the "retention of SBF" session. (**b**) IP CNO does not disrupt SBF when PVN neurons express ChR2 (instead of hM4Di). (**c**) Blue light during exposure to the CS does not significantly decrease freezing when oxytocinergic PVN neurons express hM4Di. *$p<0.05$, **$p<0.01$. Conventions as in Fig. 1. "n.s." two tailed student t-test, $p=0.8985$). For all experiments: n=3-5 animals (as indicated by the dots). Insets on left show virus injections sites and concomitant treatments. See also[1].

**Supplementary Figure 5: *In vitro* and *in vivo* electrophysiological recordings in PVN and CeA neurons demonstrating their opto,- and chemogenetic modulation**

**(a)** virus injection of an AAV expressing double floxed DIO mCherry-hM4Di under the OT promoter (pOT) in the PVN, and CAV2 expressing CRE was injected in the CeA. This combined genetic – retrograde labeling approach ensures hM4Di expression specifically in oxytocinergic neurons in the PVN that project to the CeA. Inset below shows PVN neurons expressing the AAV virus revealed by antibody against mCherry (in red) and overlap with oxytocin expression (in green). Traces below show an example of an *in vitro* electrophysiological recording of a mCherry-fluorescent hM4Di expressing neuron in slices of the PVN. Its activation by hyperpolarizing followed by a depolarizing current protocol (from -100 to +300 pA in 10 steps of 40 pAs) is decreased following perfusion of CNO thus demonstrating the functional expression of hM4Di in the OT-ergic neurons. This demonstrates that, functionally, OT neurons are indeed silenced by CNO leading to less OT release during DREADD suppression. Inset at the bottom shows the staining of OTergic fibers in the CeA (mCherry, by viral infection with this viral construct) and oxytocin receptors (by RNAscope) **(b)** virus injection of an AAV expressing double floxed DIO mCherry-ChR2 under the OT promoter (pOT) in the PVN, and CAV2 expressing CRE injected in the CeA. This combined genetic – retrograde labeling approach ensures ChR2 expression specifically in oxytocinergic neurons in the PVN that project to the CeA. Inset below shows PVN neurons expressing this mCherry-ChR2 construct as revealed by antibody against mCherry (in red) and overlap with oxytocin expression (in green). Inset below shows the staining of OTergic fibers in the CeA (mCherry, by viral infection with this viral construct) and oxytocin receptors (by RNAscope). Traces below show functional expression in *in vitro* electrophysiological slice recording of a neuron in the medial part of the central amygdala (CeM). These neurons receive inhibitory projections from OT-receptor expressing neurons in the lateral part of the CeA (CeL)[2]. Blue-light induces endogenous release of oxytocin in the CeL that activates these GABAergic neurons producing a transient increase in GABAergic currents in the CeM neuron (upper trace) that are partially blocked by simultaneous application of the oxytocin receptor antagonist OTA (middle trace) and fully blocked by additional application of CNQX (lower trace) leaving only spontaneous inhibitory postsynaptic currents. This demonstrates *in vitro* the functional expression of our construct, similar to as we have previously shown[3].

**Supplementary Figure 6. *In vivo* recording protocol and data analysis pipeline demonstrates the stability across different behavioral paradigms.**

**(a)**. The standard protocol of *in vivo* optrode recording from implantation to the end of experiments (from day 0 to day3 or day4). With little modification from behavior protocol in Figure 1a to preserve the data quality of electrophysiology **(b)** Example of optrodes with 32 channel Omnetic connectors (in blue boxes) as seen implanted in a rat head. Metal mesh was installed around the implanted electrodes as a mini-Faraday cage to decrease the electrical noise (blue square). **(c)** Upper panel: Example of online spike acquisition from Plexon system. Single unit spike waveforms were isolated by time amplitude window discrimination with a threshold 1.5 times over the baseline noise and template matching using a multichannel acquisition processor system (RASPUTIN software, Plexon). By selecting a reference channel (in which no no-spike was detected, only noise), the cross-channel artifacts and noise were eliminated. Lower panel: We aligned the spike waveforms global minima (with a scope of 100 microseconds around minima, maximum shift set to 20) and manually chose PCA features (PC1, PC2, PC3, slice 1, slice 2, timesteps etc.) to examine more densely clustered waveforms (as a same unit). We visualized spatially separated waveforms as different units in principal component 3D feature projections (as shown in Supplementary Fig. 6c, lower panel). A group of waveforms was considered to originate from a single neuron if it was defined as a discrete cluster in principal component space that was distinct from clusters for other units and if it displayed a clear refractory period (1.2 ms) in auto-correlograms (60, 61). Template waveforms were then calculated for well-separated clusters and stored for further analysis in MATLAB, as well as to track neurons over time. To ensure that the same neuron was recorded over multiple sessions (6 hours or more), we quantified the squared Mahalanobis distance, discarding neurons with unstable values. For further confirmation, we also measured cluster stability across recording sessions using the J3 and Davies-Bouldin statistics (60, 61).and PCA (lower panels) on a single tetrode in the amygdala. Automatic sorting, such as valley seeking or T-Dist E-M Scan in Plexon Sorter, was done initially just for reference. **(d1-d3)** Examples of activity of two different *in vivo* recorded neurons over a three days behavioral protocol. (Top) 2D cluster shows two units separated through PCA analysis; the unit clusters are stable from Day 1 to Day 3. (Bottom) Color code waveforms of the two units (100 traces with their average). PC1EL1, principal component electrode 1; PC1EL4, principal component electrode 4. **(e1-e3)** Autocorrelation histogram of the two neurons in d1-d3, indicating the refractory period of units are larger than 2 ms from day1 to day3. **(f1-f3)** Examples of the three neurons waveform tracking across all data section in 3-4 days, each column with plotted waveform showing a time bin of 10 min recording, which remain constant in f1 and f2 but gradually transformed in f3. Therefore f1-f2 were consider stable units and f3 an unstable unit that was removed from analysis. **(g)** Electrophysiological fingerprints of all the units recorded in CeA, demonstrating that most units exhibit a Full Width at Half Maximum (FWHM) of less than 150 microseconds, thereby classifying as putative interneurons[4]. (n=122 units recorded from N=8 rats). **(h)** The units loss among 8 rats over a 5 days recording section, number of total clear sorted units dropping from 122 in the beginning to 45 in the end.

**Supplementary Figure 7. Pharmacological and chemogenetic modulation of blue light responses in "buffer" and "fear" neurons in the CeA**

(**a1**) Experimental set-up for virus injections to render PVN OT neurons sensitive to blue light and to CNO: double virus injection of an AAV expressing double floxed DIO mCherry-ChR2 under the OT promoter (pOT) and an AAV expressing double floxed DIO mCherry-hM4Di under the OT promoter (pOT) in the PVN, and CAV2 expressing CRE injected in the CeA. This combined genetic – retrograde labeling approach ensures ChR2 and hM4Di expression specifically in oxytocinergic neurons in the PVN that project to the CeA. (**a2**) Summary of averaged z-scores of "Buffer" and **(a3)** "Fear" neuronal spiking responses to blue light (BL), blue light in the presence of OTA (BL-OTA) and blue light in the presence of CNO (BL-CNO). **(b&c)** Raster plots (top) and peri-event time histograms (bottom) BL response patterns of **(b)** a representative "Buffer" neuron and **(c)** a representative "Fear" neuron in the CeA. The raster plot represents the spikes appearing 0.5 s before and 1 s after 16 blue light exposures (at a frequency of 30 Hz during 0.5 s). **(b1)** "Buffer" neuron shows a strong excitatory response to BL, (**b2**) that is partially inhibited after IP administration of OTA and (**b3**) fully inhibited after IP administration of CNO. **(c1)** "Fear" neurons show an inhibitory response to BL **(c2)** that is partially inhibited after IP administration of OTA and **(c3)** fully inhibited after IP administration of CNO. (one way ANOVA paired tests for all with Geisser-Greenhouse correction, multiple comparisons using Tukey statistical hypothesis testing, *,**,***,**** $p<0.05$, 0.01, 0.001,0.0001, respectively; ns, not significant. a1. $F(1.838, 12.86) = 28.76$ $P<0.0001$ a2 , $F(2.336, 21.02) = 42.42$ $P<0.0001$)

**Supplementary Figure 8: Individual CS responses of Type 1 (Buffer) and Type 2 (Fear) neurons in the CeA and freezing levels across consecutive sessions of the social buffering paradigm**.

(**a**) Responses of individual CeA "Buffer" neuron to CS1 and CS2 across consecutive sessions of the social buffering paradigm as indicated by color shaded panels. Each dot and connecting line (in different colors) indicates single neuron response, violin bars indicate the distribution of cells in each session. Note: "Buffer" neurons responses to CS1 (**a1**) show a significant change across different sessions (D1-D4), but "no obvious" changes (except a decrease from Day1 to Day2) to the CS2 (**a2,** see also Figure 6). $F\ (3.556, 69.69) = 18.35$, $p < 0.0001$ in a1, $F\ (2.728, 60.01) = 5.941$, $p=0.0018$ in a2 **(b)** Responses of individual CeA "Fear" neuron to CS1 (**b1**) and CS2 (**b2)** across consecutive sessions of the social buffering paradigm. Note: the "Fear neurons" sharply increased responses to CS1 and CS2 on Day2 followed by a steep decrease selectively to CS1 during SBF and later sections. $F\ (2.701, 56.18) = 14.41$ $p < 0.0001$ in b1, $F\ (3.027, 63.57) = 11.24$,$p<0.0001$ in b2) **(c)** corresponding individual freezing responses of the recorded rats across consecutive sessions to CS1 (**c1**) and CS2 (**c2**). $F\ (0.7665, 3.679) = 9.058$, $p=0.0464$ in c1, $F\ (0.05040, 0.2268) = 16.62$ p=n.s. in **c2** All statistic test results are listed in the top right tables (one way ANOVA mixed-effects analysis for all with Geisser-Greenhouse correction, multiple comparisons using Dunnett statistical hypothesis testing). (inserted tables represent t-tests corresponding to the different colored panels ,*,**,***,**** $p$<0.05, 0.01, 0.001,0.0001, respectively; ns, not significant. ).

**Supplementary Fig 9. Fear conditioning without SBF protocol shows no signs of single unit extinction in 3 consecutive days.**

We recorded single unit responses in 3 rats that had undergone fear conditioning and "Fear recall" without subsequent SBF exposure which was replaced by only 4 times CS1 tones exposures ("No SBF"). The CeA "Fear neurons" were identified by their increased single unit firing responses after fear conditioning on Day 2 ("Fear recall"). a). Raster (top) and frequency (bottom) plot example of single unit firing response to CS (5 kHz) (0.5sec, 5Hz, red shade) across different sessions as in Figure 6. b. Individual and average z-scores of 8 neurons, recorded in 3 rats showing, upon extra exposure to the CS without SBF ("no SBF"), no significant decrease signs of extinguished responses to the CS the next days "Day 3" nor on "Day 4" consistent with maintained responses to the CS without signs of extinction. Significance was only detected between "habituation" and "fear recall" (after fear conditioning. (*$p<0.05$, n=8 cells, background colors as in Fig. 6). C) ANOVA multi comparison test with Tuke multiple comparisons, of data in b.

**Table 1 | Brain regions targeted by OTergic projections from the PVN.** OTergic neurons project throughout the brain, including to regions involved in social behavior and fear. By targeting these projections by optogenetic and chemogenetic means we found that modulation of the OT system across the brain had similar effects on freezing in our behavioral paradigm as modulation of OT signaling in the CeA directly (compare Fig. 1b with Fig. 3). This suggests that OT signaling in the CeA is key to SBF, although contributions of other brain regions cannot be excluded. The table is based on our previous characterization of OT-positive projections from the PVN[3].

**Table 2 | Information on the viruses used in the present study**

| Virus | Volume Injected | Titer | Origin |
|---|---|---|---|
| rAAV-pOT-hM4D(Gi)-mCherry (Serotype 1/2) | 500nl | ~1x10e10 vg/ml | Donation V. Grinevich (catalog :A181) |
| rAAV-pOT-ChR2-mCherry (Serotype 1/2) | 500nl | ~1x10e10 vg/ml | Donation V. Grinevich (catalog :A89) |
| rAAV-pOT-DIO-hM4D(Gi)-mCherry (Serotype 1/2) | 300nl | ~1x10e10 vg/ml | Donation V. Grinevich (catalog :A261) |
| rAAV-pOT-DIO-ChR2-mCherry (Serotype 1/2) | 300nl | 3,86x10e10 vg/ml | Donation V. Grinevich (catalog :A188) |
| CAV2-CRE (Serotype 2) | 500nl | Delivery at 6.9x10 e12 vg/ml Diluted to 2~3 x10e12 vg/ml | Montpellier vectorology platform(PVM), Biocampus Montpellier,. France |

* Virus Genome copies per ml vg/ml, which also written in physical particles pp

* each time PVM may deliver different virus titres, which need different dilution ratio. E.g. last time we ordered 2.85 dose of 2.5x10e12 pp about 50ul, and PVM delivered 40ul of 6.9x10e12 vg/ml

### References


1. Grund, T. *et al.* Chemogenetic activation of oxytocin neurons: Temporal dynamics, hormonal release, and behavioral consequences. *Psychoneuroendocrinology* **106**, 77–84 (2019).

2. Huber, D., Veinante, P. & R. Stoop. Vasopressin and oxytocin excite distinct neuronal populations in the central amygdala. *Science* **308**, 245–8 (2005).
3. Knobloch, H. S. *et al.* Evoked axonal oxytocin release in the central amygdala attenuates fear response. *Neuron* **73**, 553–566 (2012).
4. Blot, A. *et al.* Time-invariant feed-forward inhibition of Purkinje cells in the cerebellar cortex *in vivo*: Interneurons *in vivo*. *J Physiol* **594**, 2729–2749 (2016).
