## Supplementary figures and images for "Social buffering switches fear to safety encoding by oxytocin recruitment of central amygdala “buffer neurons”"

### supplementary figure 1

#### Supplementary figure 1

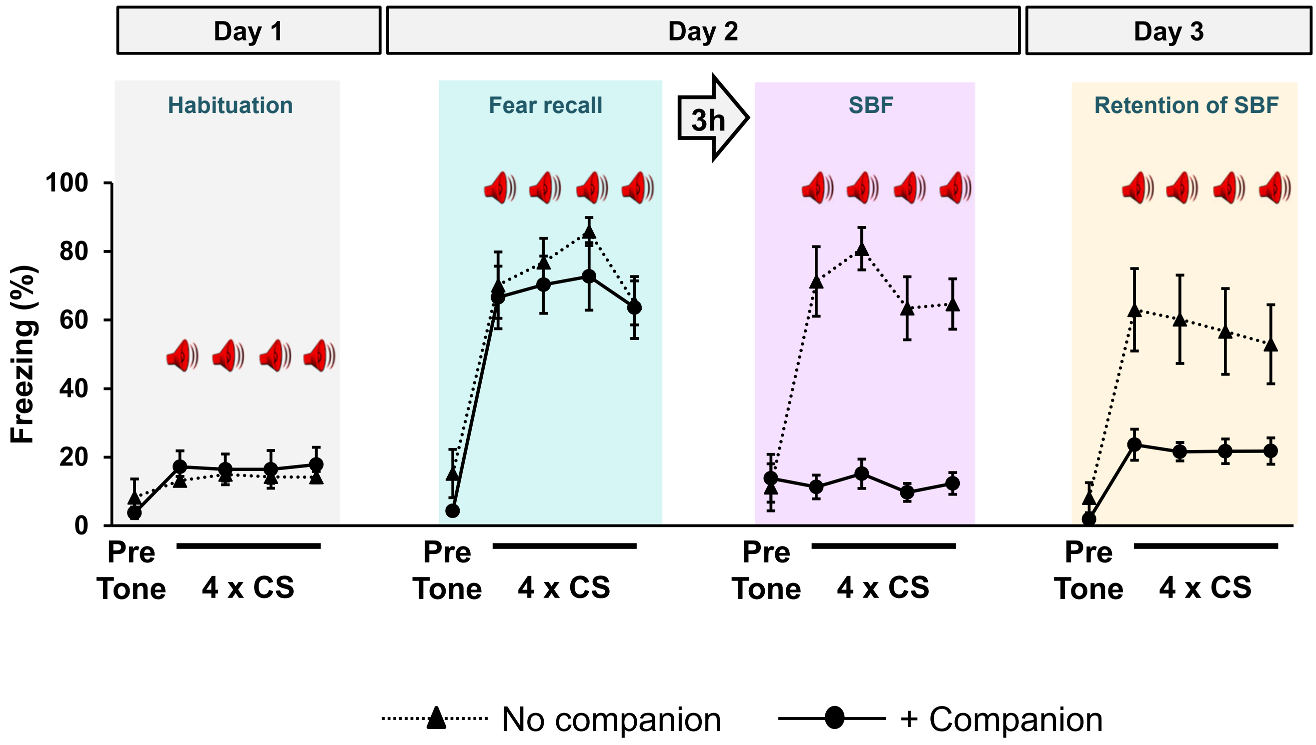

### supplementary figure 2

#### Supplementary figure 2

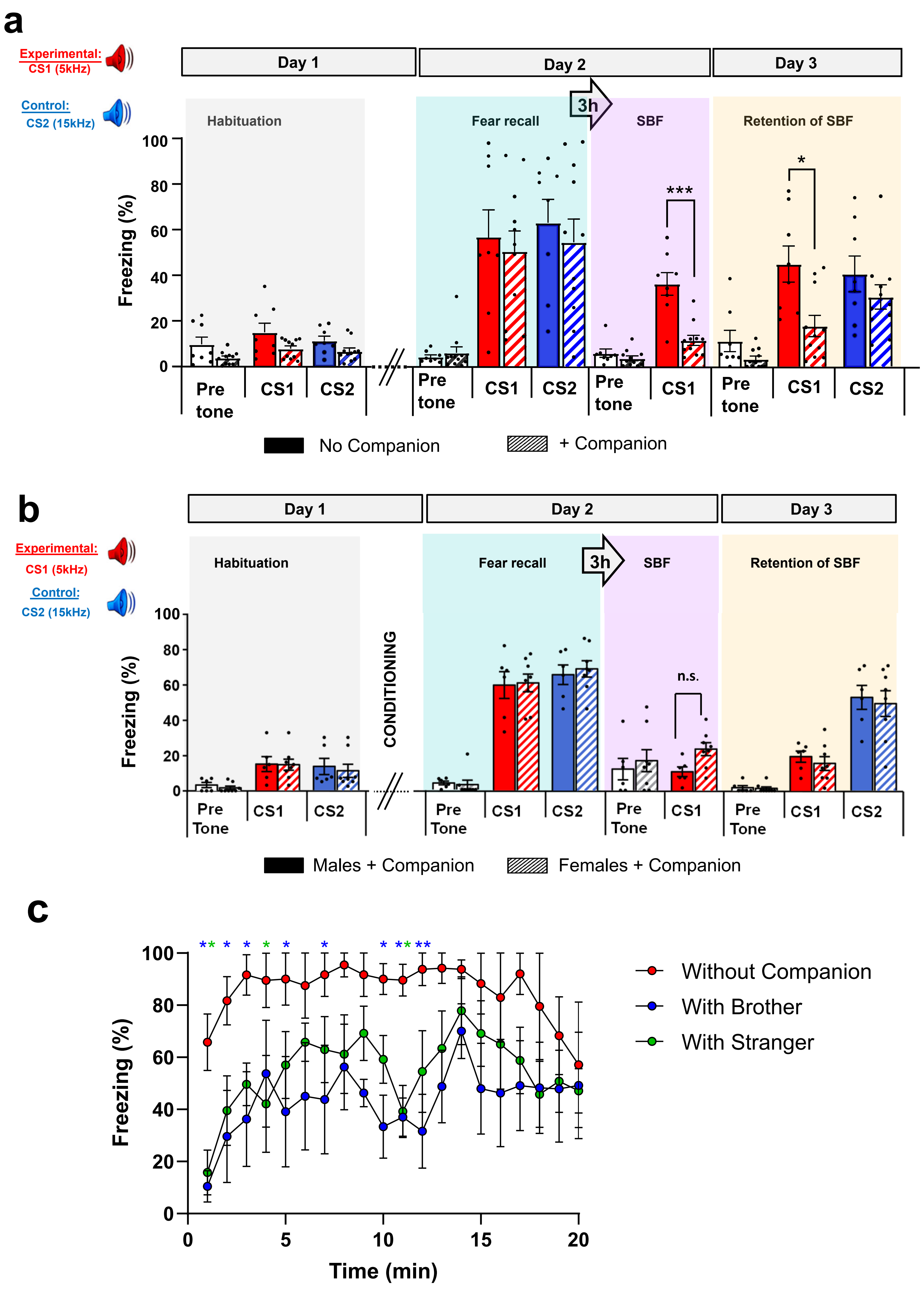

### supplementary figure 3

#### Supplementary figure 3

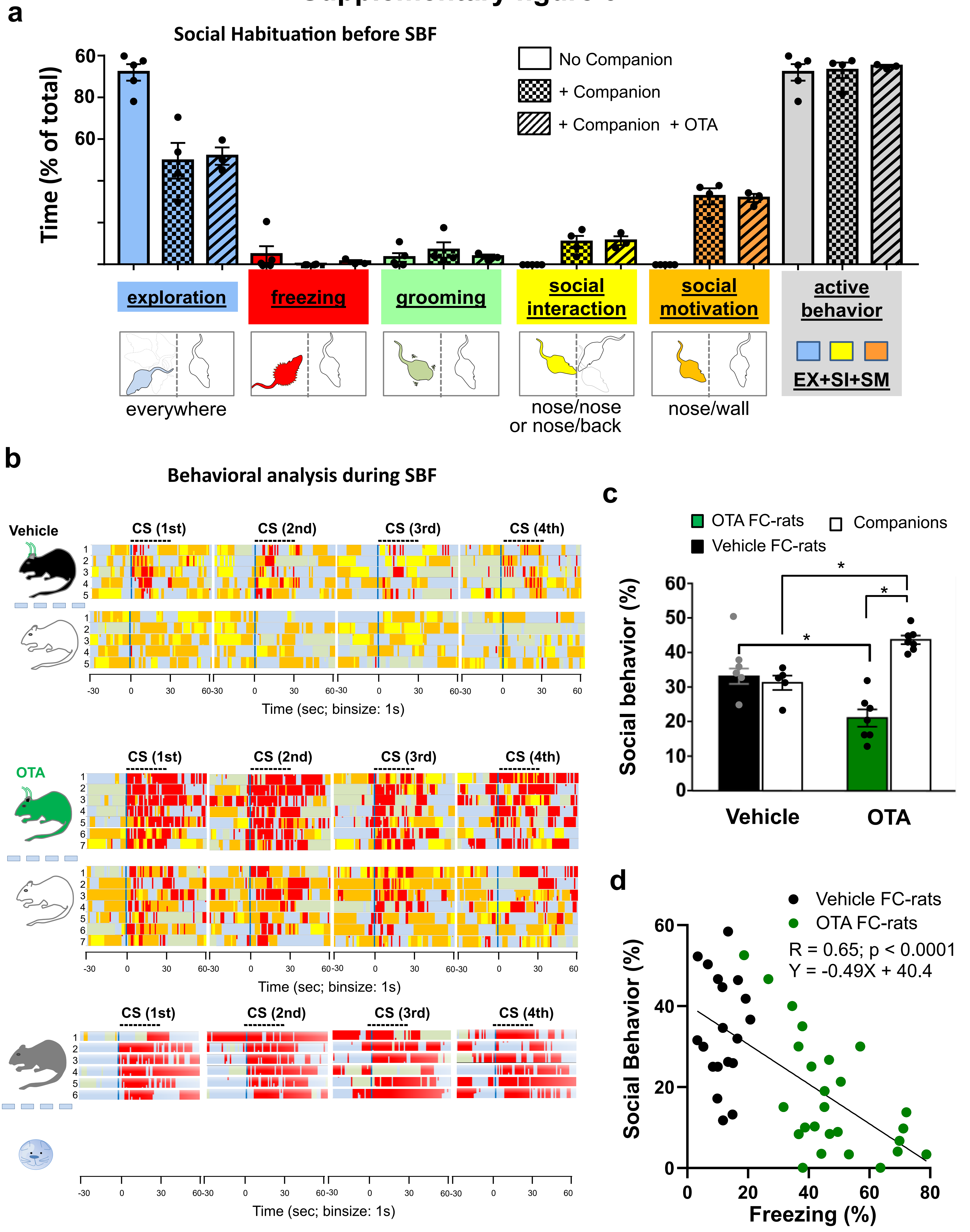

### supplementary figure 4

#### Supplementary Figure 4

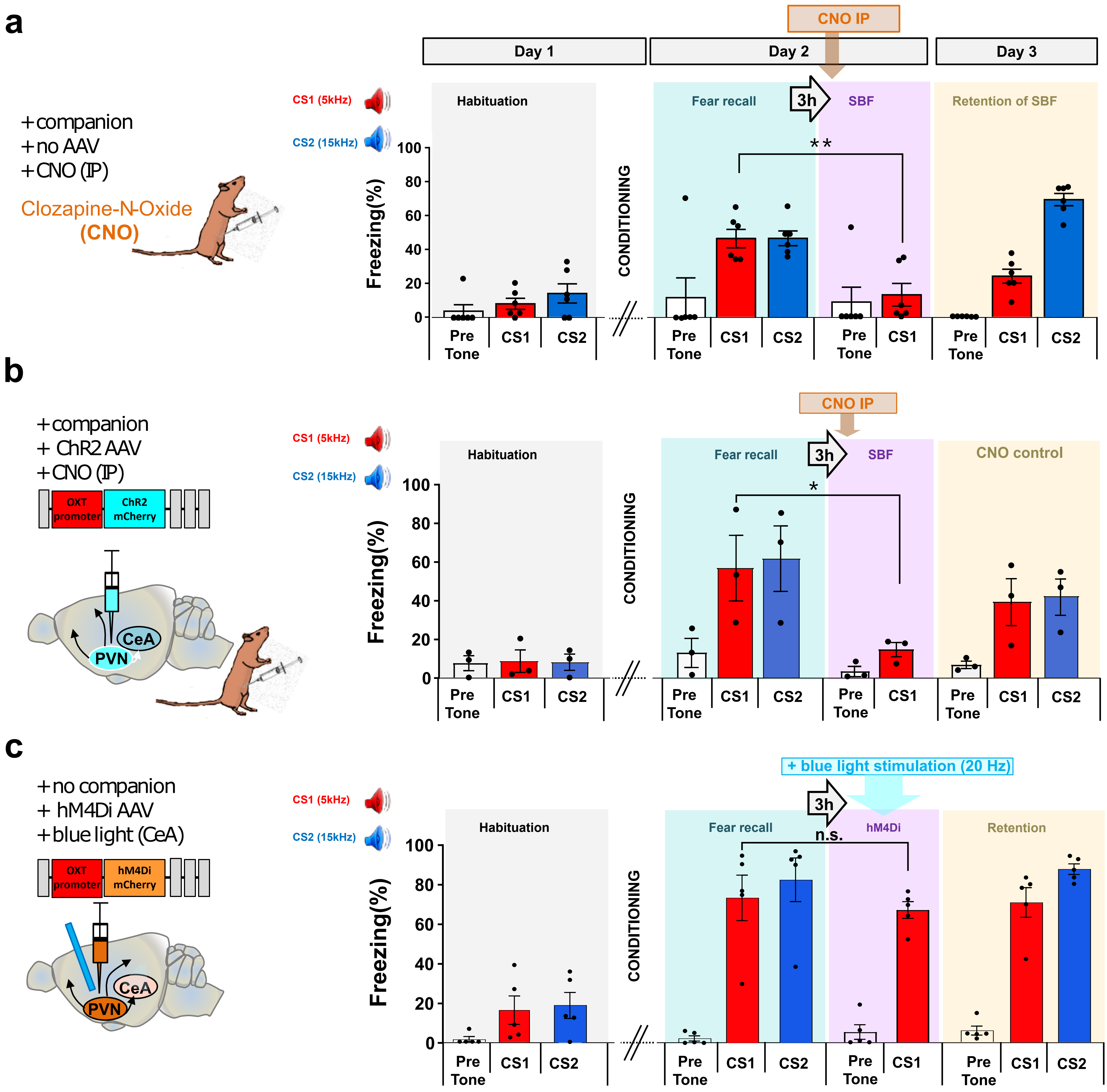

### supplementary figure 5

#### Supplementary Figure 5

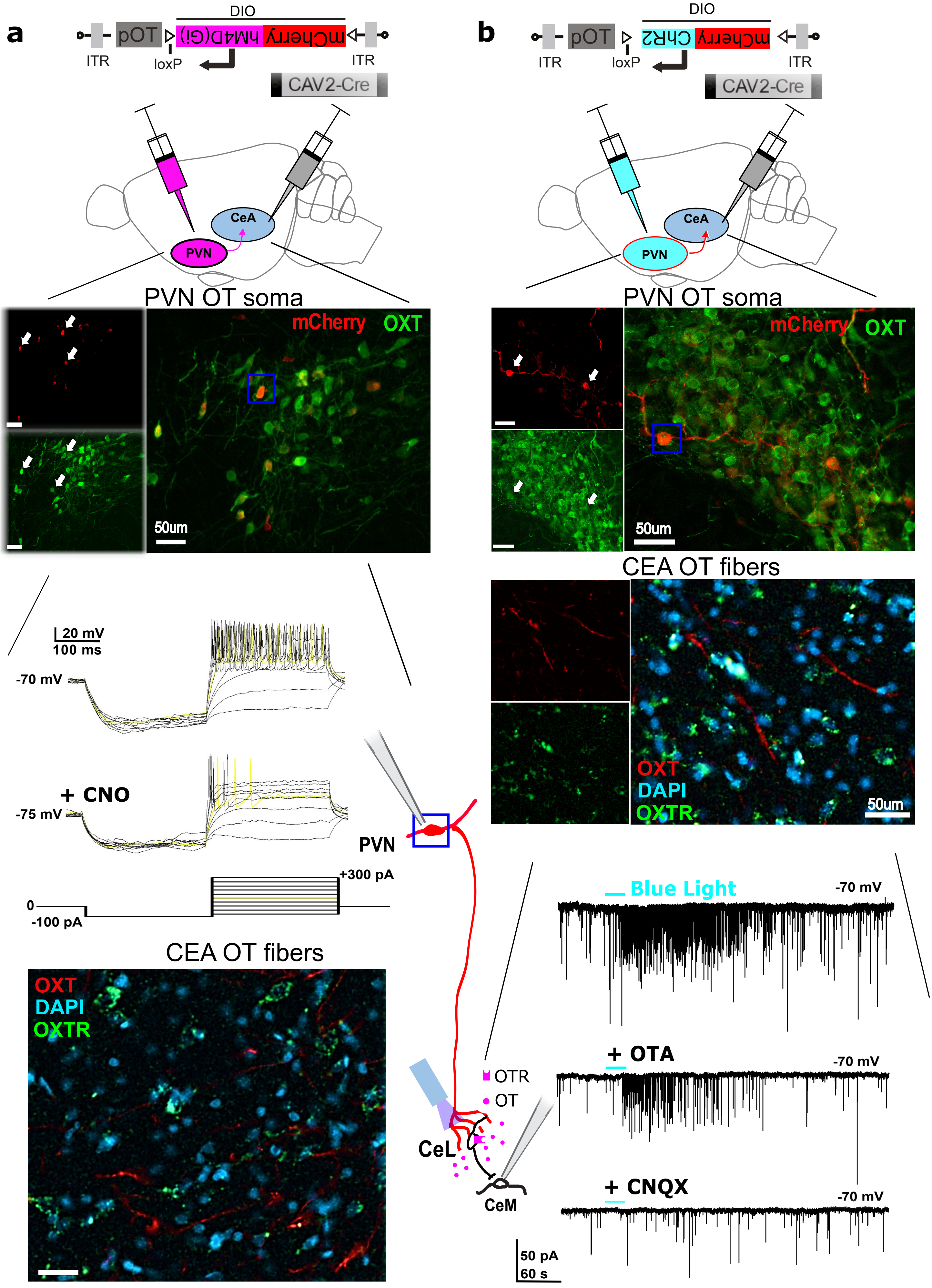

### supplementary figure 6

#### Supplementary Figure 6

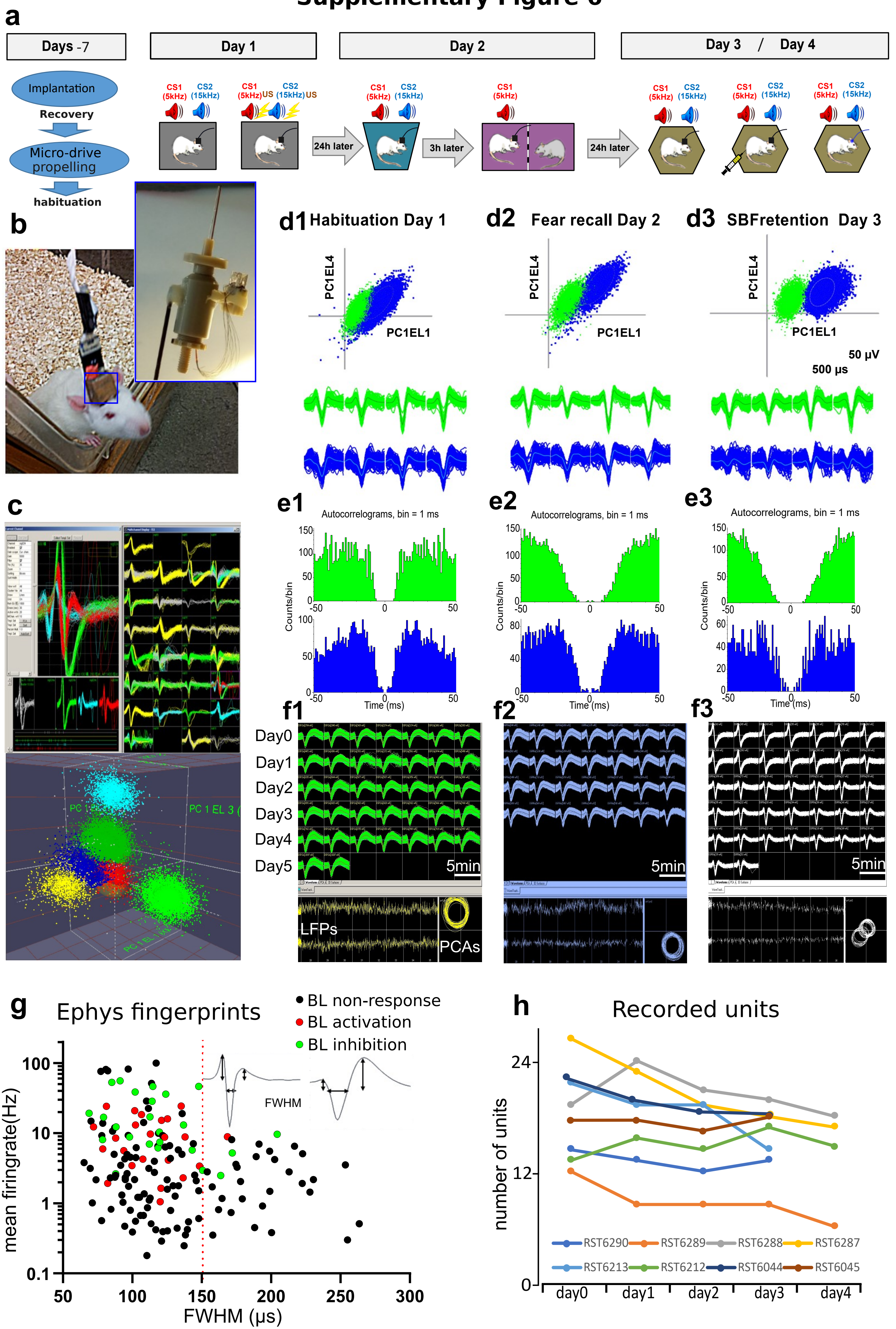

### supplementary figure 7

#### Supplementary Figure 7

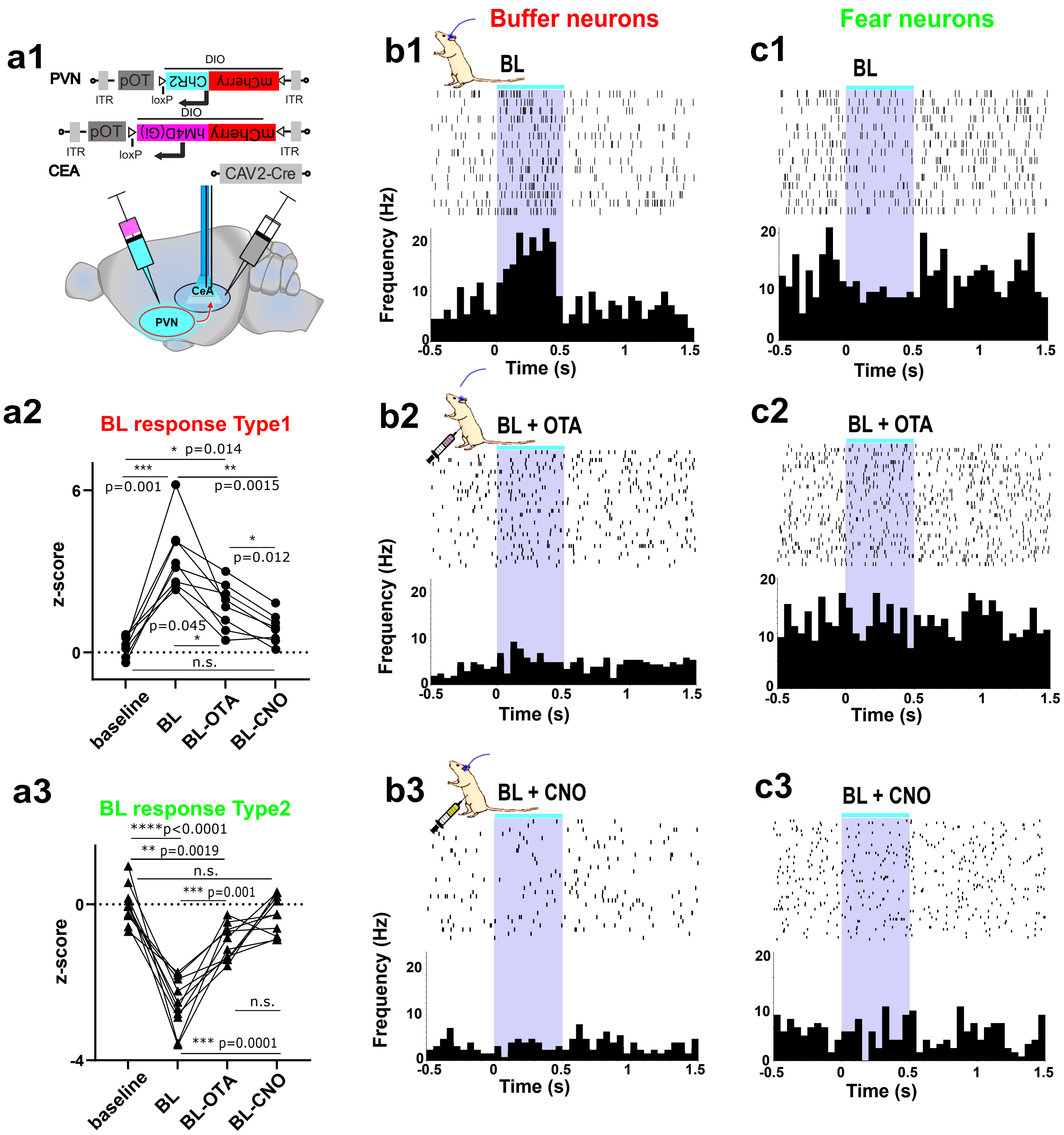

### supplementary figure 8

#### Supplementary Figure 8

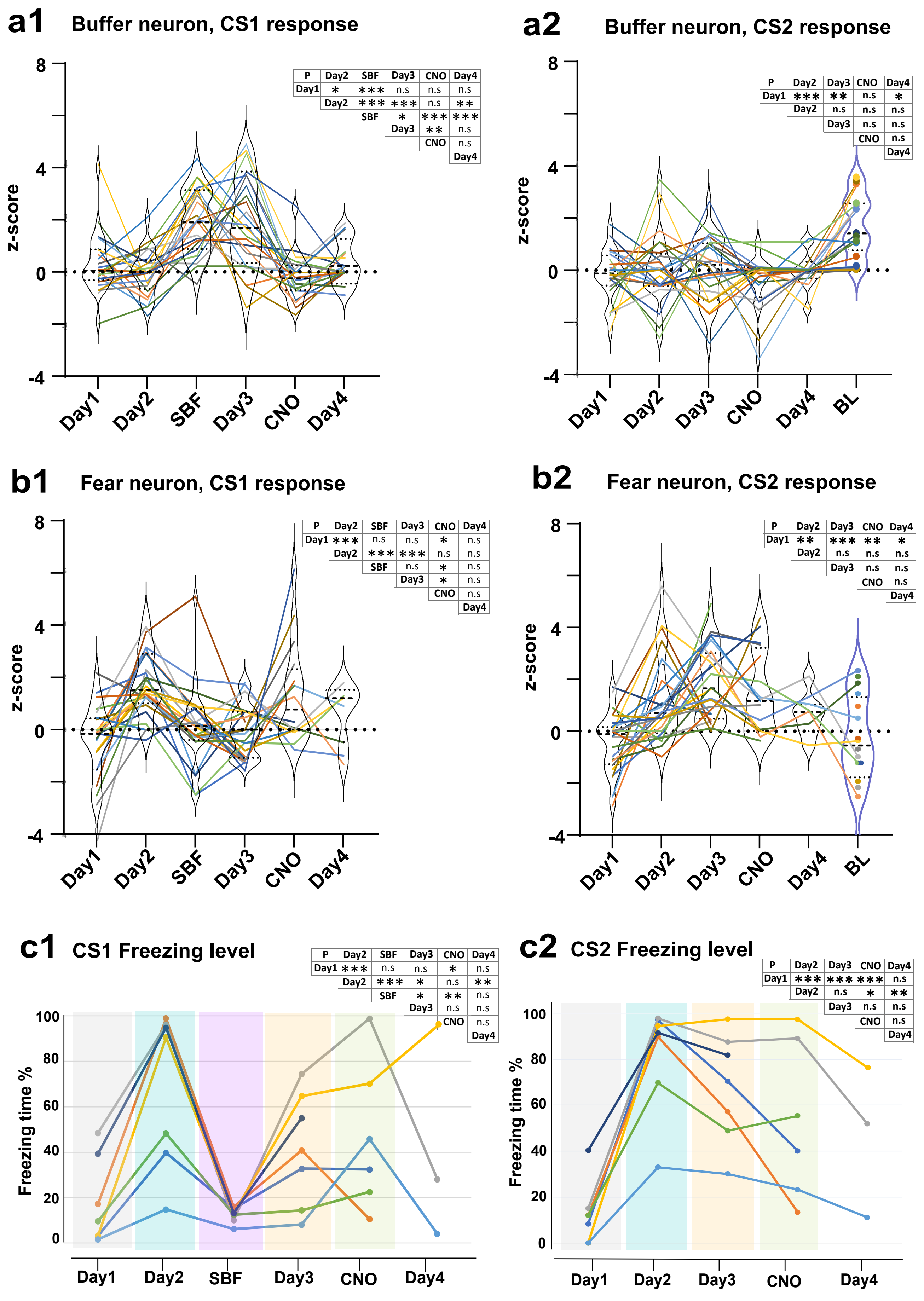
