## supplementary figure 9 for "Social buffering switches fear to safety encoding by oxytocin recruitment of central amygdala “buffer neurons”"

### Supplementory figure 9

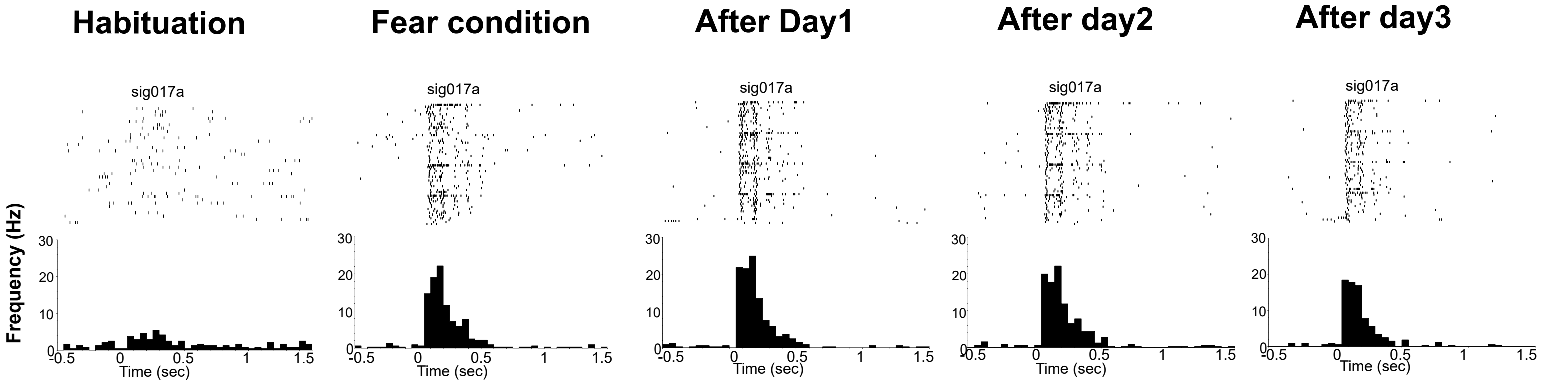

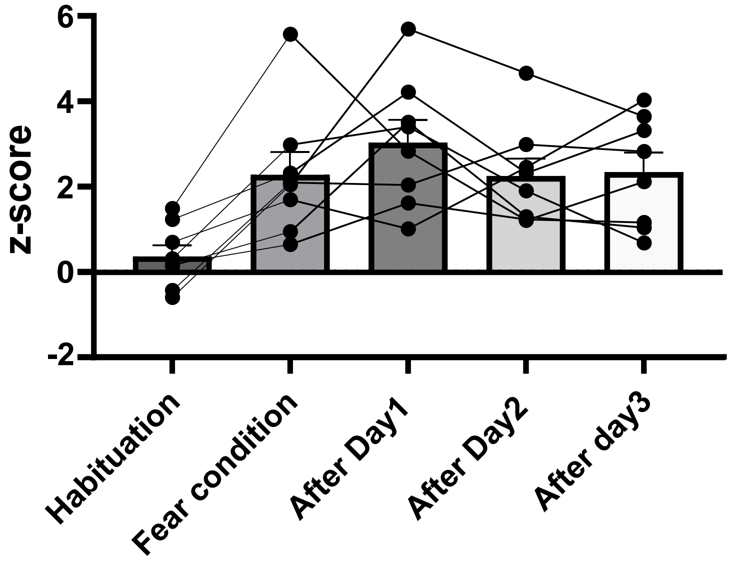

| paired t-test | Statistic: | P-value: | significanc |
|---|---|---|---|
| Habituation vs Fear condition | -4.29873 | 0.003572 | ** |
| Habituation vs After Day1 | -4.17653 | 0.004155 | ** |
| Habituation vs After Day2 | -3.11812 | 0.016888 | * |
| Habituation vs After day3 | -3.79327 | 0.006774 | ** |
| Fear condition vs After Day1 | -1.0623 | 0.32337 | n.s. |
| Fear condition vs After Day2 | 0.043317 | 0.966659 | n.s. |
| Fear condition vs After day3 | -0.09167 | 0.929527 | n.s. |
| After Day1 vs After Day2 | 1.643156 | 0.144352 | n.s. |
| After Day1 vs After day3 | 1.025902 | 0.339077 | n.s. |
| After Day2 vs After day3 | -0.2711 | 0.794134 | n.s. |

| Repeated measures ANOVA summary | | | | Was the matching effective? | |
|---|---|---|---|---|---|
| Assume sphericity? | No | | | F | 1.771 |
| F | 5.606 | | | P value | 0.1331 |
| P value | 0.009 | | | P value summary | ns |
| P value summary | ** | | | Is there significant match | No |
| Statistically significant (P < 0.05)? | Yes | | | R squared | 0.1973 |
| Geisser-Greenhouse's epsilon | 0.639 | | | | |
| R squared | 0.445 | | | | |
| ANOVA table | SS | DF | MS | F (DFn, DFd) | P value |
| Treatment (between columns) | 31.75 | 4 | 7.9 | F (2.558, 17.90) = 5.606 | P=0.0089 |
| Individual (between rows) | 17.55 | 7 | 2.5 | F (7, 28) = 1.771 | P=0.1331 |
| Residual (random) | 39.64 | 28 | 1.4 | | |
| Total | 88.94 | 39 | | | |
